# Phenotypic and molecular characterization of extended spectrum β-lactamase producing *Escherichia coli* and *Klebsiella pneumoniae* isolates from various samples of animal origin from Assam, India

**DOI:** 10.1101/2020.05.28.122705

**Authors:** Leena Das, Probodh Borah, R.K. Sharma, Dipika Malakar, G.K. Saikia, K. Sharma, S. Tamuly, Rupam Dutta

## Abstract

Extended-spectrum beta-lactamase (ESBL) producing Enterobacteriaceae has become a major threat globally. Here we have characterized ESBL producing *E. coli* and *K. pneumoniae* from various sources, studied antibiogram and resistance gene profiles. Out of 385 samples, 31 (8.05%) were positive for ESBL producing *E. coli*. Such isolates could be recovered from 10.05, 8.33, 15.63, 6.67 and 4.35 per cent of cattle milk, curd, chicken, pork and cattle faeces samples, respectively. A total of 59 (15.32%) samples were positive for ESBL producing *K. pneumoniae*, which were isolated from 14.35, 6.25, 21.43 and 34.78 per cent cattle milk, chicken, beef and cattle faeces, respectively. All the 90 isolates were confirmed as ESBL producers by CDT and ESBL-E strip tests. Antibiogram revealed that 74.19% and 69.49% of the ESBL producing *E. coli* and *K. pneumoniae* isolates, respectively showed resistance to ceftizoxime, 25.81% and 23.73% to both co-trimoxazole and tetracycline, 19.35% and 25.42% to ciprofloxacin, 9.68% and 16.95% to chloramphenicol, 3.23% and 5.08% to pipercillin-tazobactam, and 3.23% and 3.39% to gentamicin. Resistance gene profiling showed *bla*CTX-M gene as most predominant (100%). The *bla*TEM gene was found in 54.84% and 55.93%, *bla*SHV gene in 90.32% and 77.97%, *Sul* 1 gene in 90.32% and 86.44% of ESBL producing *E. coli* and *K. pneumoniae* isolates, respectively. The *Int*1 gene was detected in 70.97% and 62.71% isolates, while *qnr*B gene was found in 3.23% and 10.17% of *E. coli* and K*. pneumoniae* isolates, respectively.

## Introduction

Antibiotic resistance is a major problem of public health concern. Among the multi-drug resistant pathogens, the extended-spectrum beta-lactamase producing Enterobacteriaceae have emerged as a threat for human health worldwide (29). Extended spectrum β-lactamases (ESBLs) are enzymes carried by plasmids that show resistance to 3^rd^ and 4^th^ generation cephalosporins and also to monobactam group of drugs. However, ESBLs are still sensitive to carbapenems and cephamycins. ESBLs are most commonly detected in Enterobacteriaceae like *Klebsiella pneumoniae* and *Escherichia coli*. These enzymes can be inhibited by β-lactamase inhibitors like clavulanic acid, sulbactam and tazobactam (28). These extended spectrum β-lactamases include TEM and SHV β-lactamase families, whereas other β-lactamases such as CTX-M, PER and KPC have also been described recently (7). Majority of ESBLs in clinical isolates have been identified as SHV or TEM types, which have evolved from narrow-spectrum β-lactamases such as TEM-1, −2 and SHV-1.4. The most prevalent and commonly found beta-lactamase is TEM-1. It has been estimated that due to presence of TEM-1, more than 90% ampicillin resistance occurs in *E. coli* (22). For detection of ESBL in Enterobacteriaceae, particularly for *E. coli, Klebsiella* spp. and *Proteus* spp., the Clinical and Laboratory Standards Institute (CLSI; formerly known as the National Committee for Clinical Laboratory Standards) (5) and the Health Protection Agency (HPA) in UK suggested some guidelines (3). As per these guidelines, ESBL producing organisms can be detected by supplementation of either 8 mg/L (CLSI) or 1 mg/L (HPA) of cefpodoxime, 1 mg/L each of cefotaxime, ceftazidime, ceftriaxone, or aztreonam in media followed by phenotypical confirmation using both cefotaxime and ceftazidime in combination with clavulanic acid and by use of ESBL E-strips. Several genetic mechanisms have been found to be involved in transmission of antimicrobial resistance genes. The term “mobilome” (24, 36) is used for various mobile genetic elements (MGEs) including plasmids, transposons (Tn), insertion sequences (IS), integrons (intI) and introns. Integrons are DNA fragments embedded in gene cassettes capable of disseminating antimicrobial resistance genes using mobile genetic elements (35). Class 1 (*intI1*) and class 2 (*intI2*) integrons are most commonly involved in the antibiotic resistance mechanism (11, 33, 23, 26, 16). The present study attempted to ascertain the prevalence of ESBL producing *E. coli* and *K. pneumoniae* among domestic animals and to determine their antibiogram and resistance gene profiles.

## Materials and methods

### Reference strains

In the present study, *Escherichia coli* ATCC^®^25922 and *Klebsiella pneumoniae* ATCC^®^700603 were used as an ESBL negative and ESBL positive controls, respectively.

### Sampling

A total of 385 samples were collected from different districts of Assam, *viz*. Kamrup, Nagaon, Morigaon, Barpeta and Bongaigaon during the period from May, 2017 to June 2018, and were examined for the presence of ESBL-producing *Escherichia coli* and *Klebsiella pneumoniae*. The samples included 209 cattle milk, 23 goat milk, 12 curd (indigenously prepared), 32 chicken meat, 15 pork, 14 beef, and faecal samples of cattle (69) and goat (11).

### Isolation and identification of suspected ESBL producing *E. coli* and *K. pneumoniae*

For primary isolation, the samples were inoculated in 5 ml of Luria Burtoni broth (Himedia, India) supplemented with cefotaxime @ 1 mg/L and the tubes were incubated aerobically at 37° C for 24 hours. A loopful of broth culture was then streaked onto Mac Conkey’s Lactose Agar (MLA) (Himedia, India) plates supplemented with cefotaxime @ 1 mg/L and incubated for a period of 24 hours at 37° C. Subsequently, the suspected colonies of ESBL producing ESBL producing *E. coli* colonies were subcultured on Eosin Methylene Blue Agar (EMBA) and *Klebsiella* spp. were subcultured on MLA plates (Himedia, India) for purification. The plates were then incubated aerobically at 37° C for 24 hours. Characterization and preliminary identification of the suspected isolates of *Klebsiella* spp. (27) and *E. coli* (18, 2) were done on the basis of morphology and colony characteristics. Biochemical tests were performed for identification of *Klebsiella pneumoniae* (12, 8) and *E.coli* (18) isolates. Molecular confirmation of the suspected ESBL producing *Klebsiella* spp. and *E. coli* isolates was done by PCR amplification of *rpo*B and *uid*A genes, respectively. DNA extraction was done using Bacterial DNA Isolation Kit (Geneaid, Taiwan). The Primer sequences are described in Table 1. The cycling conditions for *uid*A and *rpo*B gene were as follows: initial denaturation at 95°C for 5 min, 35 cycles of each steps including denaturation at 95°C for 30 sec, annealing at 54°C for 1 min for *uid*A and 51°C for 1 min for *rpo*B gene and elongation at 72°C for 30 sec, followed by final elongation at 72°C for 5 min.

**Table 1:**
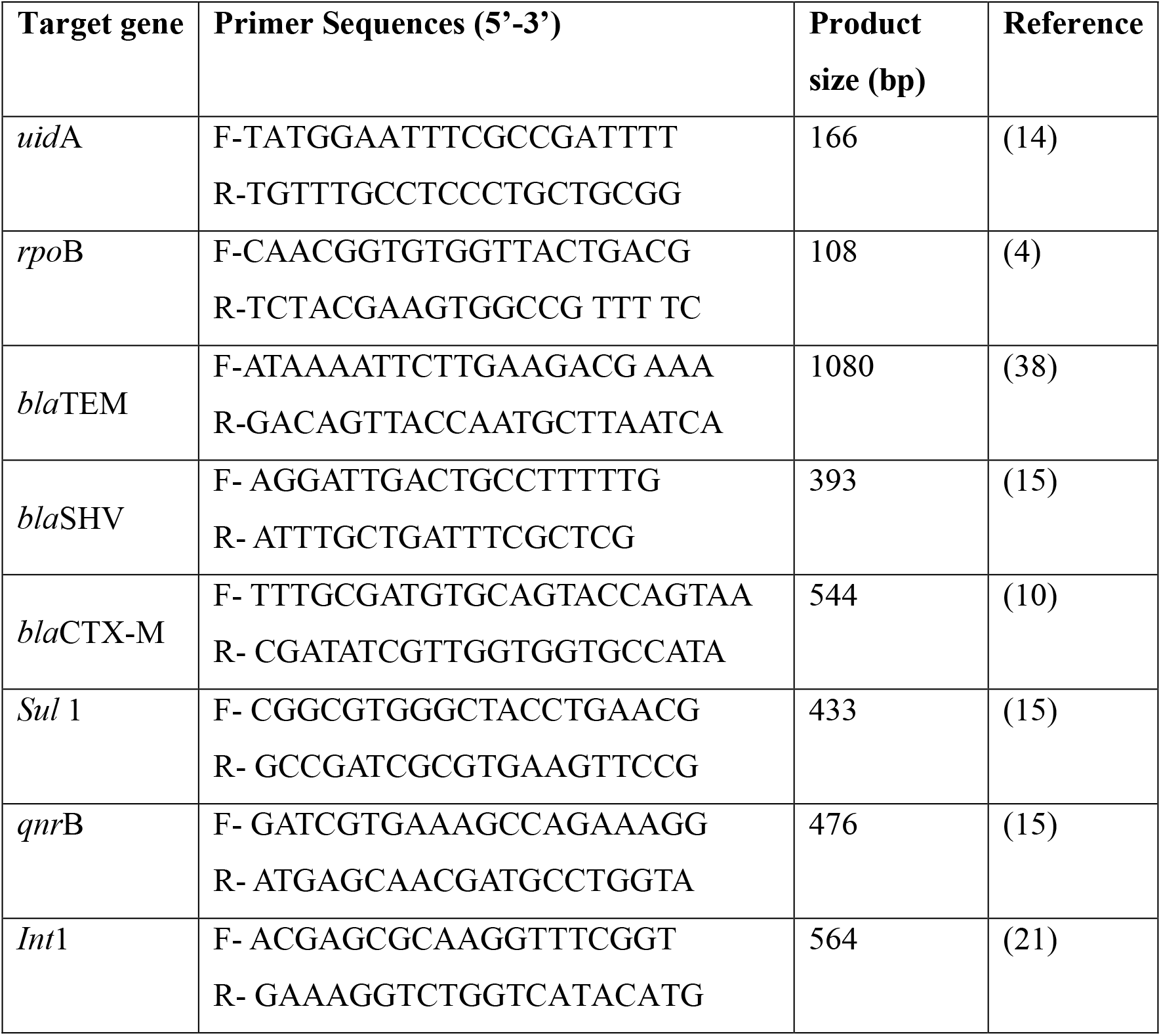
PRIMER SEQUENCES FOR DETECTION DIFFERENT GENE BY SIMPLEX PCR USED IN THIS STUDY.

### Phenotypic confirmation of ESBL producing *E. coli* and *K. pneumoniae*

Confirmation of the suspected ESBL isolates was done by three different antibiotic tests *viz*. antibiotic sensitivity assay, Combination of Disc Diffusion Tests (CDT) and ESBL-E-test according to the Clinical and Laboratory Standards Institute (6) guidelines. All the suspected ESBL isolates were first subjected to drug sensitivity assay against six different antibiotics, *viz*. cefotaxime 30 μg, ceftriaxone 30 μg, cefpodoxime 10 μg, ceftazidime 30 μg, cefepime 30 μg and aztreonam 30 μg (Himedia, India). In the Combination of Disc Diffusion Tests (CDT), discs containing a third generation antibiotic, cefotaxime (30 μg) and a combination of cefotaxime/clavulanic acid (30/10 μg) were used. After overnight incubation at 37°C, a difference in the diameter of zone of inhibition by ≥ 5 mm between cefotaxime 30 μg disc and cefotaxime/clavulanic acid 30/10 μg disc was interpreted as ESBL positive. Then the isolates were subjected to ESBL –E test (Himedia, India), where one side of the strip contained three antibiotics, *viz*. ceftazidime, cefotaxime and cefepime, and the other side additionally contained clavulanic acid. After overnight incubation at 37°C, if the ratio of the value of the mix to that of the mix + clavulanic acid was found to be ≥ 8, the isolate was considered as an ESBL producer.

### Detection of AmpC β- lactamase and carbapenamase production

AmpC β- lactamase production was detected by cefoxitin-cloxacillin double disc synergy (CC-DDS) test (30). The test was performed using antimicrobial discs containing a combination of cefoxitin 30 μg and cefoxitin-cloxacillin 200 μg. This test is based on the principle that AmpC production can be inhibited by addition of cloxacillin in combination with antibiotic. A zone diameter difference of ≥ 4 mm between cefoxitin 30 μg disc and cefoxitin-cloxacillin 30-200 μg disc was interpreted as AmpC positive.

Carbapenemase production was detected by Modified Hodge Test (MHT) and E-test (32). Modified-Hodge test was performed by preparing 0.5 McFarland dilution of the *E. coli* ATCC 25922 strain in 5 ml of Mueller-Hinton broth (Himedia, India). The culture was diluted 1:10 by adding 4.5 ml of Mueller-Hinton broth into 0.5 ml of broth culture comparable to the 0.5 McFarland standards. The diluted (1:10) culture was streaked on to Mueller-Hinton agar plate and an ertapenem (10 μg) antibiotic disc (Himedia, India) was placed at the centre of the plate, and the test isolates were streaked in a straight line from the edge of the disc to the edge of the plate. Appearance of clover-leaf structure after overnight incubation indicated carbapenemase production.

The isolates were then subjected E-test for detection of carbapenemase production. The E-strip (Himedia, India) containing imipenem (4-256 mcg/ml) in one side and imipenem and EDTA (1-64 mcg/ml) in combination on the other side was used for the test. After overnight incubation at 37°C, if the ratio of the value for Imipenem (IPM) to that of Imipenem + EDTA (IPM+EDTA) was more than 8 or if a zone of inhibition was observed on the side coated with Imipenem+EDTA whereas no zone on the opposite the side coated with Imipenem, culture was interpreted as carbapenemase producer.

### Antibiogram profile of ESBL producing *E. coli* and *K. pneumoniae* isolates

All the ESBL producing isolates were subjected to antibiotic sensitivity assay against nine different antibiotics, *viz*. co-trimoxazole 25 μg, chloramphenicol 30 μg, tetracycline 30 μg, gentamicin 30 μg, ceftizoxime 30 μg, ciprofloxacin 5μg, imipenem 10 μg, meropenem 10 μg, and pipercillin-tazobactam 100/10 μg, and interpretation was done as per the guidelines of Clinical and Laboratory Standards Institute (6).

### Screening of resistance genes by PCR

All the ESBL-producing *E. coli* and *K. pneumoniae* isolates were screened for the presence of six important antimicrobial resistance genes, *viz*. β-lactamase genes- *bla*TEM, *bla*SHV, *bla*CTX-M, sulfonamide resistance gene- *Sul* 1, plasmid mediated quinolone resistance gene- *qnr*B and class 1 integron-*Int*1 by PCR using the primers shown in Table 1. The cycling conditions for *bla*TEM, *bla*SHV, *bla*CTX-M, *Sul* 1, *Int*1 and *qnr*B genes were as follows: initial denaturation at 95°C for 5 min, 35 cycles for each steps of denaturation at 95°C for 30 sec, annealing at 43°C (***bla*TEM**) and 58°C (***Sul* 1**) for 1min, elongation at 72°C for 30 sec, followed by final elongation at 72°C for 5 min. Likewise, initial denaturation at 95°C for 5 min, 30 cycles for each steps of denaturation at 95°C for 30 sec, annealing at 45°C (***bla*SHV**) and at 53°C (***bla*CTX-M**) for 1min, elongation at 72°C for 30 sec, final elongation at 72°C for 5 min. For the rest two genes, *Int*1 and *qnr*B; initial denaturation at 95°C for 5 min, 35 cycles for each steps of denaturation at 95°C for 30 sec, annealing at 60°C (***qnr*B**) and at 57°C (***Int*1**) for 45 sec and elongation at 72°C for 30 sec, followed by final elongation at 72°C for 5 min.

## Results & Discussion

ESBLs have been reported globally by many researchers in organisms belonging to *Enterobacteriaceae* and *Pseudomonadaceae* families, but most commonly in *Klebsiella pneumoniae* and *Escherichia coli*. The emergence of ESBL has become an important public health concern because of association with increased morbidity, mortality and healthcare costs (29). The present investigation was conducted with an aim to detect and characterize ESBL producing *E. coli* and *K. pneumoniae* isolates from various animal sources.

Out of 385 different samples examined, 31 (8.05%) were positive for ESBL producing *E. coli* as confirmed by PCR amplification of *uid*A gene (Figure 1) and 59 (15.32%) were positive for ESBL producing *Klebsiella pneumoniae* (Figure 2) confirmed by amplification of *rpo*B gene. (Table 2). In the present study, 21 (10.05%) and 30 (14.35%) of 209 cattle raw milk samples yielded ESBL producing *E. coli* and *K. pneumoniae*, respectively, while none of the 23 goat raw milk samples yielded ESBL producing *E. coli* and *K. pneumoniae*. None of the 12 curd samples tested in the present study yielded ESBL producing *K. pneumoniae*, while only 1 (8.33%) of these samples yielded *E. coli*. A total of 5 (15.63%) and 2 (6.25%) samples among 32 chicken meat samples examined were found positive for ESBL producing *E. coli* and *K. pneumoniae*, respectively. None of the 15 pork samples yielded *K. pneumoniae* isolates, while only 1 (6.67%) yielded ESBL producing *E. coli*. No ESBL producing *E. coli* isolate could be obtained from the 14 beef samples and only 3 (21.43%) of them yielded *K. pneumoniae*. While 3 (4.35%) and 24 (34.78%) of 69 cattle faecal samples yielded ESBL producing *E. coli* and *K. pneumoniae* isolates, respectively, none of the 11 goat faecal samples were found to be positive for any such isolate. In the present study, high prevalence of ESBL producing *E. coli* was detected in meat (9.84%), followed by milk (9.05%), curd (8.33%) and faeces (3.75%). On the other hand, occurrence of ESBL producing *K. pneumoniae* was higher in faecal samples (30%), followed by milk (12.93%) and meat (8.19%) (Table 2). In conformity to the present study, ESBL producing *K. pneumoniae* were detected in 6.7% bovine milk samples in Eastern and North-Eastern India (19). Higher prevalence of ESBL isolates in milk observed in the present study might be due to smaller sample size. In another study (9), similar prevalence rate of 2%, 3.1% and 1% were found in pig, cattle and pigeon faecal samples, respectively in Hong Kong. A higher isolation rate of 35% ESBL producing *K. pneumoniae* was reported in beef and chicken samples in Turkey (13), which might be due to geographical and climatic variations or influence of managemental and host factors on acquiring infection from the environment. In this study no ESBL isolates could be obtained from milk and faecal samples of goat, which might be due to lesser use of antibiotic therapy in this species compared to cattle. No ESBL producing *E. coli* isolate could be obtained from beef samples and similarly, no ESBL producing *K. pneumoniae* isolate could be recovered from pork and curd samples.

**FIG. 1.**
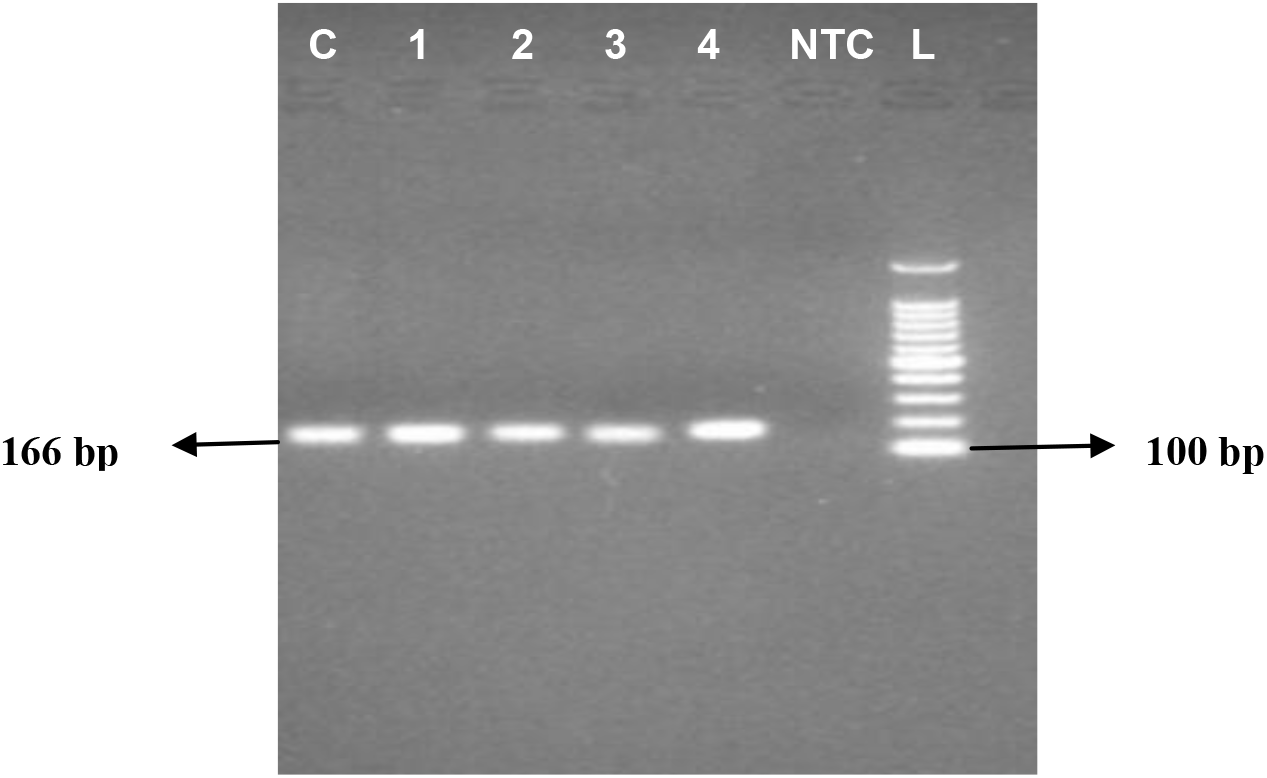
AMPLIFIED PRODUCTS OF *uid*A GENE IN 1.5% AGAROSE GEL FOR CONFIRMATION OF *E.coli* ISOLATES. Lane 1: Positive control, Lane 2 to 5: Amplified products, Lane 6: No-template control, Lane 7: 100 bp DNA ladder

**FIG. 2.**
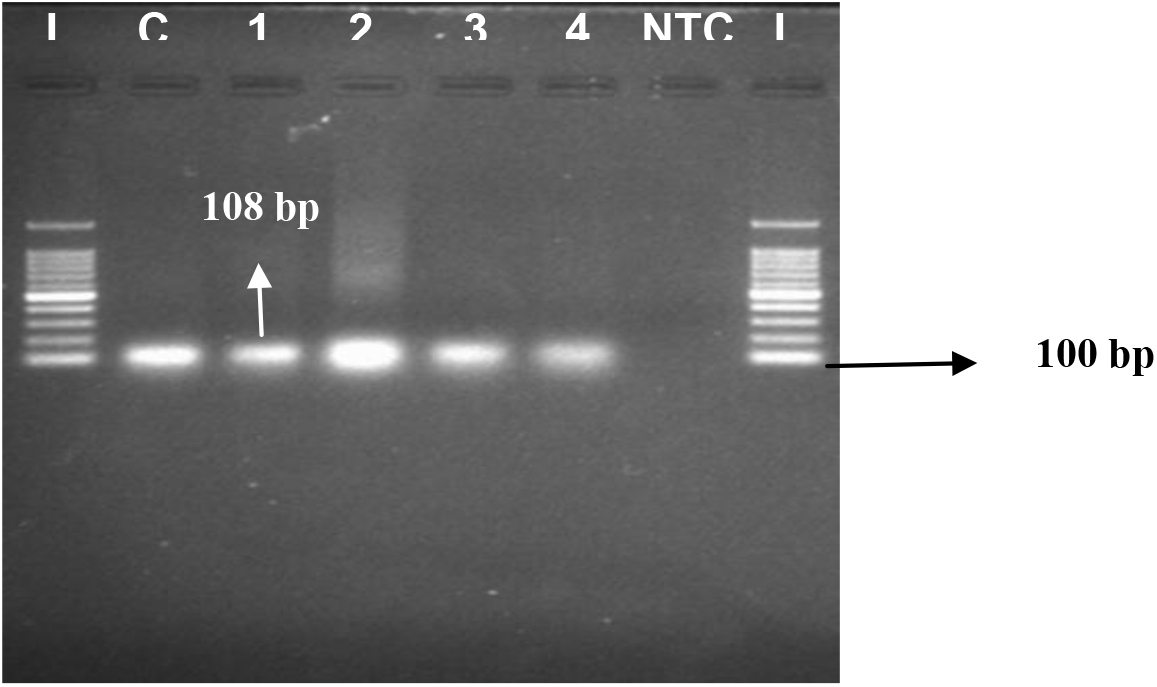
AMPLIFIED PRODUCTS OF *rpo* B GENE IN 1.5% AGAROSE GEL FOR CONFIRMATION OF *K.pneumoniae* ISOLATES. Lane 1 and 8: 100 bp DNA ladder Lane 2: Positive control, Lane 3 to 6: Amplified products, Lane 7: No-template control,

**Table 2:** ISOLATION OF ESBL PRODUCING *Escherichia coli* AND *Klebsiella pneumoniae* FROM VARIOUS SOURCES.

| Nature of sample | Source | Number of samples tested | Number positive for ESBL isolates |  |  |  |
| --- | --- | --- | --- | --- | --- | --- |
|  |  |  | *E. coli* |  | *K. pneumoniae* |  |
| Raw milk | Cattle | 209 | 21 (10.05) | **21(9.05)** | 30 (14.35) | **30(12.93)** |
|  | Goat | 23 | - |  | - |  |
| Curd (Indigenous) | Cattle | 12 | 1 (8.33) | **1 (8.33)** | - | - |
| Meat | Chicken | 32 | 5 (15.63) | **6(9.84)** | 2 (6.25) | **5(8.19)** |
|  | Pork | 15 | 1 (6.67) |  | - |  |
|  | Beef | 14 | - |  | 3 (21.43) |  |
| Faeces | Cattle | 69 | 3 (4.35) | **3(3.75)** | 24 (34.78) | **24(30.0)** |
|  | Goat | 11 | - |  | - |  |
| **Total** |  | **385** | **31 (8.05)** |  | **59 (15.32)** |  |

Figures in parenthesis indicate percentages

In the initial antibiotic sensitivity assay, all the 31 *E. coli* isolates showed 100% resistance to ceftriaxone and cefpodoxime, followed by cefotaxime (96.77%), aztreonam (83.87%), cefepime (77.42%) and ceftazidime (58.06%). All the 59 *K. pneumoniae* (100%) isolates showed resistance to cefpodoxime, followed by cefotaxime (96.61%) and ceftriaxone (93.22%), aztreonam (89.83%), ceftazidime (77.97%) and cefepime (50.85%). All the 90 isolates (31 *E. coli* and 59 *K. pneumoniae* together) were confirmed as ESBL producer by CDT and ESBL E-tests. Drug sensitivity assay of ESBL producing *E. coli* isolates showed the highest resistance to ceftizoxime (74.19%), followed co-trimoxazole and tetracycline (25.81% each) and ciprofloxacin (19.35%). The isolates showed least resistance to chloramphenicol (9.68%), gentamicin and pipercillin-tazobactam (3.23% each). The ESBL producing *K. pneumoniae* isolates showed the highest level of resistance to ceftizoxime (69.49%), followed by ciprofloxacin (25.42%), and co-trimoxazole and tetracycline (23.73% each). Comparatively lower resistance was shown to chloramphenicol (16.95%), pipercillin-tazobactam (5.08%) and gentamicin (3.39%). None of the 90 ESBL isolates showed resistance against imipenem and meropenem. Out of 31 ESBL producing *E. coli* isolates, 6.45%, 9.68%, 3.23% and 9.68% showed resistance to five, four, three and two antimicrobial agents, respectively. On the other hand, 5.08%, 20.34% and 25.42% of the ESBL producing *K. pneumoniae* (59) isolates showed resistance to four, three and two different antimicrobial agents, respectively. Overall, 29.32% of *E. coli* and 50.85% of *K. pneumoniae* isolates showed multiple drug resistance. These findings were in partial agreement with a study from Pondicherry, India (37), who found higher sensitivity of *E. coli* and *K. pneumoniae* to imipenen (100% and 98%), pipercillin-tazobactam (84% and 68%) and amikacin (68% and 40%) respectively. Another study from Nigeria reported 14% and 16 % sensitivity ESBL producing *E. coli* and *K. pneumoniae* isolates, respectively to amoxicillin/clavulanate and 43% and 21%, respectively to ceftazidime. Most bacterial pathogens showed more than 60% sensitivity to ceftriaxone, gentamicin and levofloxacin (25).

None of the 90 ESBL producing isolates were found to be positive for AmpC production by CC-DDS test and for production of carbapenemase by MHT and Imipenem+ EDTA E-strip test. ESBLs and AmpC β-lactamases together are becoming global threat to animals as well as human beings for causing multidrug resistance resulting in treatment failure. However, in the present study, no ESBL (*E. coli* and *K. pneumoniae*) isolate was found to be positive for AmpC production. This might be due to lesser possibility of co-production of AmpC and ESBL, or over-production of ESBL might lower the expression of AmpC. Likewise, no MBL (Metallo-β lactamase) producing isolate could be recovered in the present study. This indicates the possibility of treating patients affected with multi drug resistant bacteria using carbapenems.

Antibiotic resistance is reported to be mediated by many variants of β-lactamases like TEM, SHV, CTX-M, PER and KPC (7). Screening for presence of resistance genes showed *bla*CTX-M (Figure 5) gene to be present in all the 90 (100%) ESBL producing *E. coli* and *K. pneumoniae* isolates. In case of *E. coli*, 28 (90.32%) isolates were found to harbor *bla*SHV (Figure 4) and *Sul* 1 genes (Figure 6). *Int*1 gene (Figure 8) was detected in 22 (70.97 %) isolates and *bla*TEM (Figure 3) in 17 (54.84%) isolates. Only 1 (3.23%) *E. coli* isolate was found to harbor *qnr*B gene (Figure 7). In case of *K. pneumoniae*, 51 (86.44%) isolates were found to harbor *Sul* 1 gene, followed by *bla*SHV gene in 46 (77.97%) isolates, *Int*1 in 37 (62.71%); *bla*TEM in 33 (55.93%) and *qnr*B gene in 6 (10.17%) isolates. In the present study, many ESBL producing *E. coli* and *K. pneumoniae* isolates were found to carry more than one resistant genes (Table 3). One *E. coli* and six *K. pneumoniae* isolates were found to carry all the six resistant genes. All other five genes except *qnr*B were found in 11 (35.48%) *E. coli* and 19 (32.20%) *K. pneumoniae* isolates, respectively. The genes *bla*TEM, *bla*CTX-M, *Sul* 1 and *Int*1 together were found only in two *K. pneumoniae* isolates, whereas, *bla*SHV, *bla*CTX-M, *Sul* 1 and *Int*1 were found in five *E. coli* and six *K. pneumoniae* isolates. All the three *bla* genes, *viz. bla*TEM, *bla*SHV and *bla*CTX-M were found in one *K. pneumoniae* isolate. Combination of *bla*CTX-M, *Sul* 1 and *Int*1 genes were detected in two isolates each of *E. coli* and *K. pneumoniae*. Likewise, combination *bla*SHV and *bla*CTX-M was detected in two *K. pneumoniae* isolates. The present findings are in close agreement with a report from Saudi Arabia (39) where, occurrence of *bla*CTX-M and *bla*TEM genes was found to be 95.3% and 83.9% of ESBL producing *E. coli* isolates. However, *bla*OXA and *bla*SHV were detected by them only in 6.6% and 5.2% isolates, respectively, which was not in agreement with the present study. Another study conducted in French revealed presence of *bla*CTX-M, *bla*TEM and *bla*SHV genes in 39.6%, 24.6% and 3.7% *E. coli* isolates, whereas *Klebsiella* spp. less frequently harboured *bla*CTX-M (5.9%), *bla*TEM (2.7%) and *bla*SHV (1.6%) genes (20). These findings were not in conformity to that of the present study. In contrast to the present study, report from East Africa (34) showed presence of *qnr*B, *qnr*S and *qnr*A genes in 74 (47.74%), 73(47.10%) and 4(2.58%) ESBL (*E. coli* and *Klebsiella* spp.) isolates out of a total 107 isolates. The gene *qnr*B was detected in 41(38.32%) *E. coli* and 33 (30.84%) *Klebsiella* spp., *qnr*S in 48(44.86%) *E. coli* and 25 (23.36%) *Klebsiella* spp. and all the four isolates harboring *qnr*S gene were of *Klebsiella* spp. Another study (1) reported a prevalence rate of 81%, 67% and 2.29% for *Sul*1, *Sul*2 and *Sul*3 genes, respectively among *E. coli* isolates. Their findings were in close agreement with that of the present study. However on the contrary, *Int*1 gene was detected in 11.76% *E. coli* isolates (17). Similarly, another study reported a prevalence rate of 44% for *Int*1 gene, followed by 6% for *Int*2 gene among *K. pneumoniae* isolates (31), which were in partial agreement with the present findings.

**FIG. 3.**
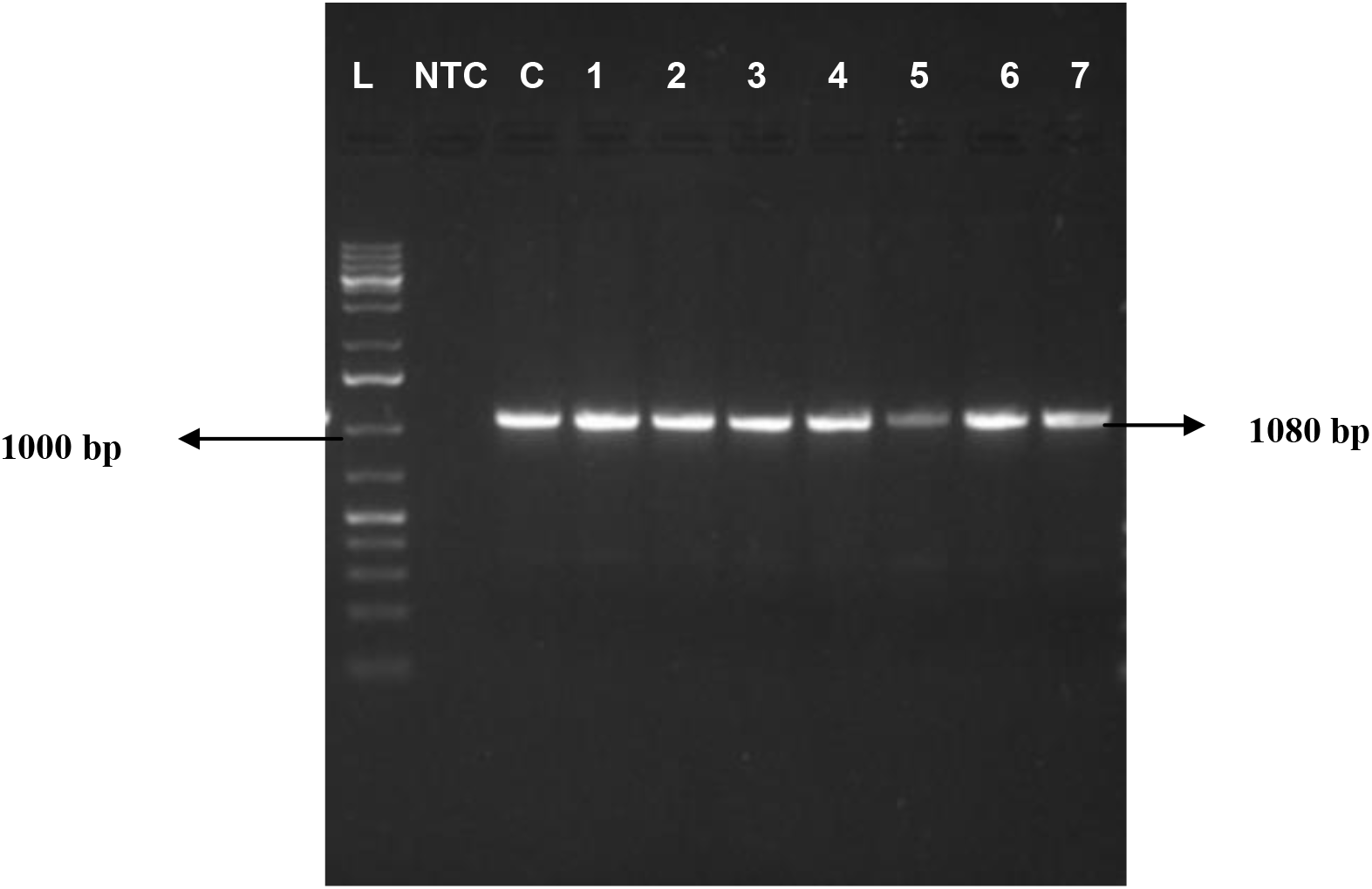
AMPLIFIED PRODUCTS OF *bla*TEM GENE IN 1.5% AGAROSE GEL. Lane 1: 1Kb DNA ladder Lane 2: No-template control, Lane 3: Positive control Lane 4 to 6: Amplified products of *E. coli* Lane 7 to 10: Amplified products of *K. pneumoniae*

**FIG. 4.**
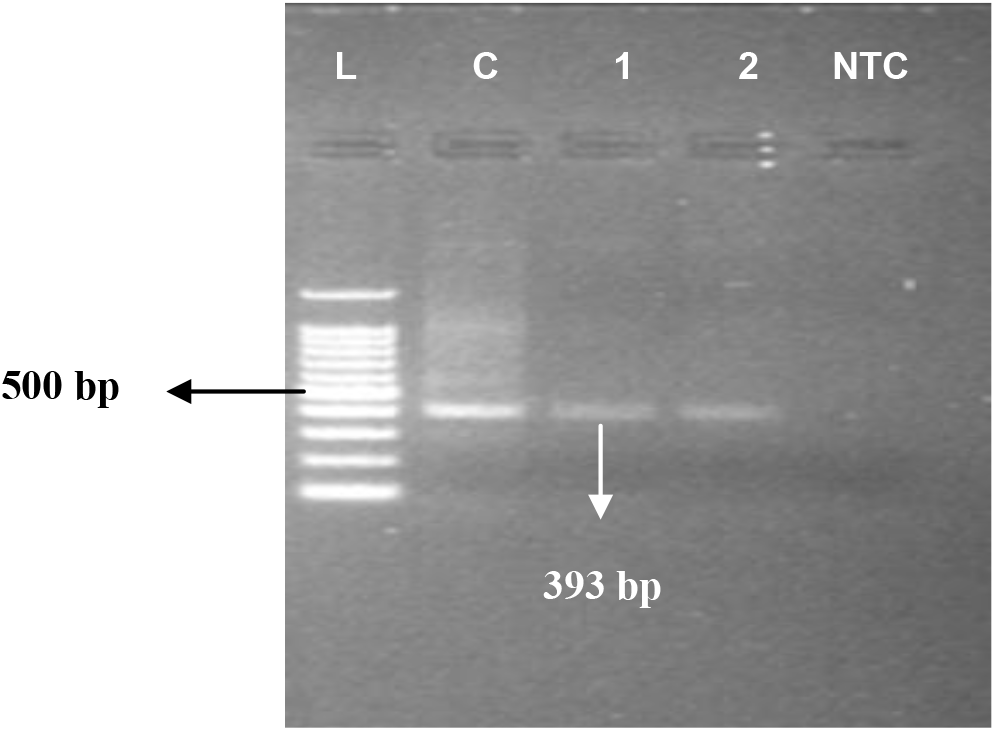
AMPLIFIED PRODUCTS OF *bla*SHV GENE IN 1.5% AGAROSE GEL. Lane 1: 100 bp DNA ladder Lane 2: Positive control Lane 3: Amplified product of *E. coli* Lane 4: Amplified products of *K. pneumoniae* Lane 5: No-template control

**FIG. 5.**
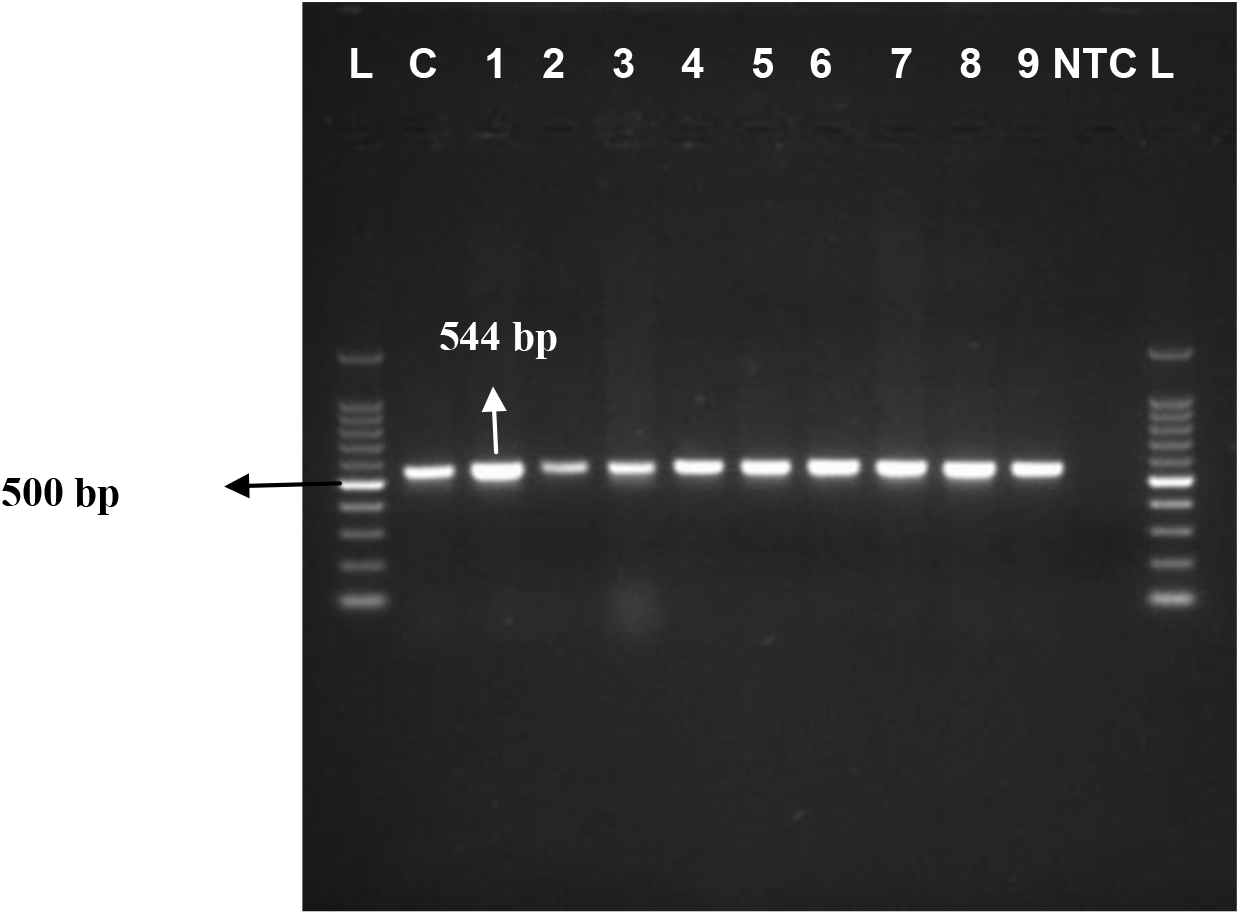
AMPLIFIED PRODUCTS OF *bla*CTX-M GENE IN 1.5% AGAROSE GEL. Lane 1 and 13: 100 bp DNA ladder Lane 2: Positive control Lane 3 to 6: Amplified products of *E. coli* Lane 7 to11: Amplified products of *K. pneumoniae* Lane 12: No-template control

**FIG. 6.**
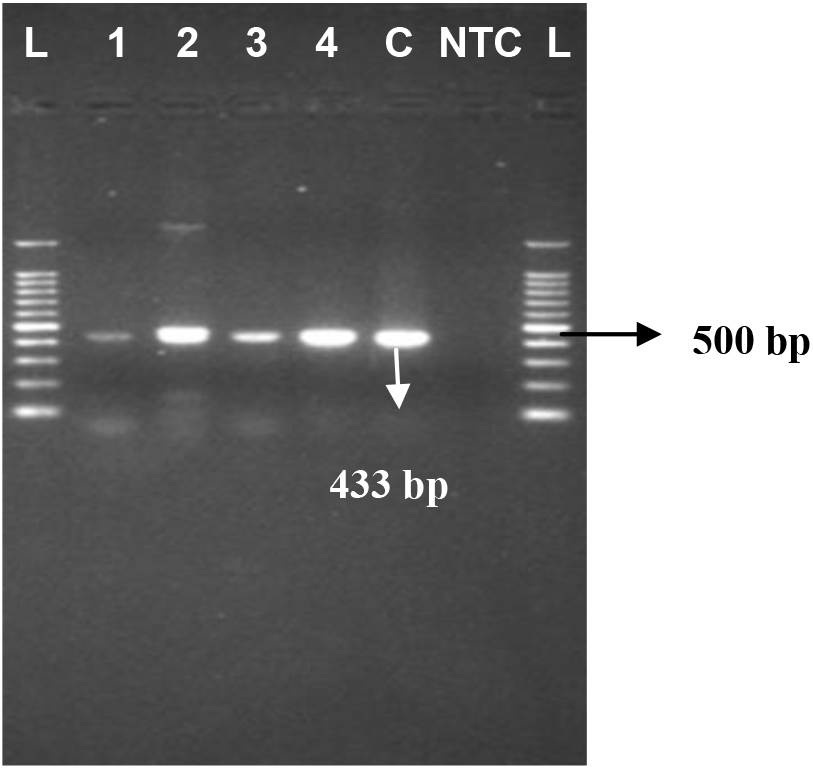
AMPLIFIED PRODUCTS OF *Sul*1 GENE IN 1.5% AGAROSE GEL. Lane 1 and 8: 100 bp DNA ladder, Lane 2 and 3: Amplified products of *E. coli*, Lane 4 and 5: Amplified products of *K. pneumoniae* Lane 6: Positive control; Lane 7: No-template control

**FIG. 7.**
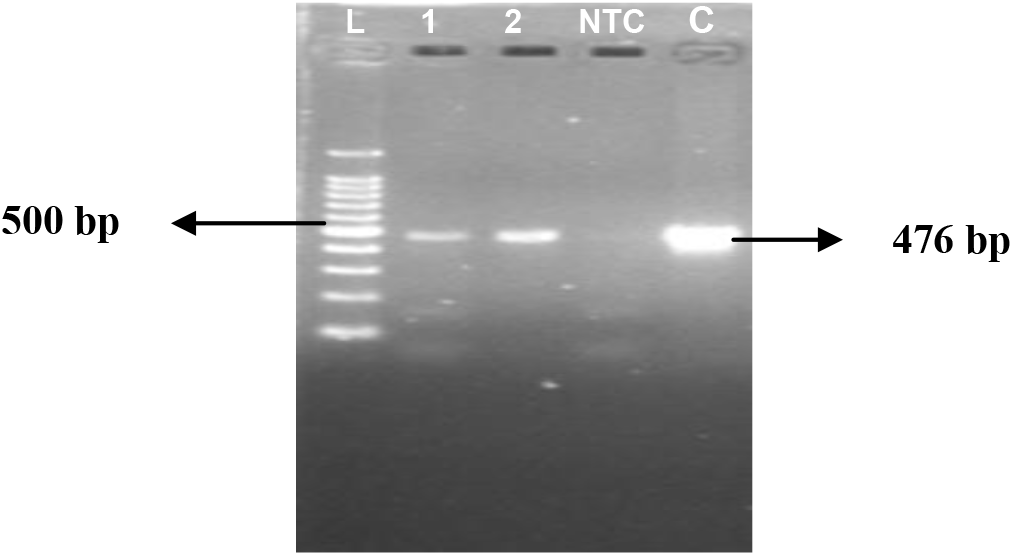
AMPLIFIED PRODUCTS OF *qnr*B GENE IN 1.5% AGAROSE GEL. Lane 1: 100 bp DNA ladder, Lane 2: Amplified product of *E. coli*; Lane 3: Amplified product *K. pneumoniae*, Lane 4: No-template control, Lane 5: Positive control

**FIG. 8.**
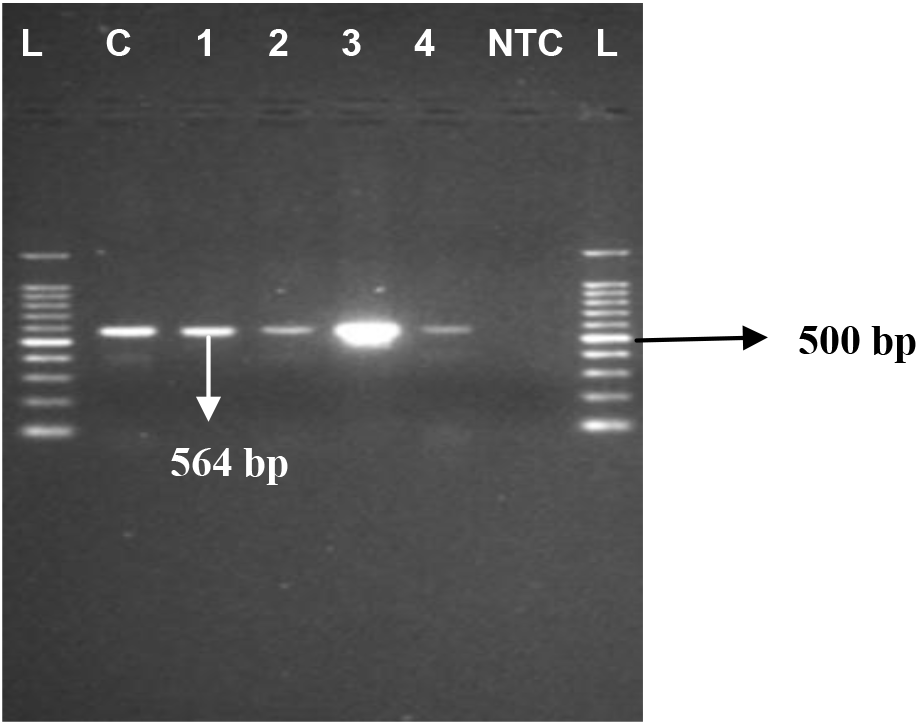
AMPLIFIED PRODUCTS OF *Int* 1 GENE IN 1.5% AGAROSE GEL. Lane 1 and 8: 100 bp DNA ladder, Lane 2: Positive control, Lane 3 and 4: Amplified products of *E. coli*, Lane 5 and 6: Amplified products of *K. pneumoniae*, Lane 7: No-template control

**Table 3:** CHARACTERIZATION OF ESBL PRODUCING *E.coli* AND *K.pneumoniae* ISOLATES GENOTYPICALLY.

| Combination of genes | No.of *E.coli* isolates | No.of*K.pneumoniae* isolates |
| --- | --- | --- |
| *bla*TEM, *bla*SHV, *bla*CTX-M, *Sul* 1, *qnr*B, *Int*1 | 1 | 6 |
| *bla*TEM, *bla*SHV, *bla*CTX-M, *Sul* 1, *Int*1 | 11 | 19 |
| *bla*TEM, *bla*CTX-M, *Sul* 1, *Int*1 | - | 2 |
| *bla*SHV, *bla*CTX-M, *Sul* 1, *Int*1 | 5 | 6 |
| *bla*TEM, *bla*SHV, *bla*CTX-M | - | 1 |
| *bla*CTX-M, *Sul* 1, *Int*1 | 2 | 2 |
| *bla*SHV, *bla*CTX-M | - | 2 |

## Conclusions

The present study provided preliminary data on prevalence of ESBL producing *E. coli* and *K. pneumoniae* in various samples of animal origin in Assam. Based on the study, it could be concluded that prevalence of ESBL producing *E. coli* and *K. pneumoniae* is still lower in the state compared to other parts of India and abroad. However, the prevalence of ESBL producing *K. pneumoniae* was higher compared to *E. coli*.

## Acknowledgments

The authors acknowledge the support received from the State Biotech Hub (Assam), College of Veterinary Science, Khanapara, Assam, India, funded by the Department of Biotechnology, Government of India and Department of Animal Biotechnology, College of Veterinary Science, Khanapara for conducting the study.

